# Implicit auditory perception of local and global irregularities in passive listening condition

**DOI:** 10.1101/2020.07.23.218925

**Authors:** Krystsina Liaukovich, Yulia Ukraintseva, Olga Martynova

## Abstract

The auditory system detects differences in sounds at an implicit level, but data on this difference might not be sufficient to make explicit discrimination. The biomarkers of implicit auditory memory of ambiguous stimuli could shed light on unconscious auditory processing and implicit auditory learning. Mismatch negativity (MMN) and P3a, components of event-related potentials (ERPs) reflecting stimuli discrimination without direct attention, were previously detected in response to the local (short-term) irregularity in the auditory sequence even in an unconscious state. At the same time, P3b was elicited only in case of direct attention in response to the global (long-term) irregularity. In this study, we applied the local-global auditory paradigm to obtain possible electrophysiological signatures of implicit detection of hardly distinguishable auditory stimuli. ERPs were recorded from 20 healthy volunteers during active discrimination of deviant sounds in the old-ball sequence and passive listening of the same sounds in the sequence with local-global irregularity. The discrimination task consisted of two blocks with different deviant sounds targeted to respond. The sound discrimination accuracy was at an average of 40 %, implying the difficulty of explicit sound recognition. Comparing ERPs to standard and deviant sounds, we found posterior negativity in ERP around 450-600 ms in response to targeted deviant sounds. MMN was significant only in response to non-target deviants. In the passive local-global paradigm, we observed an anterior positivity (284-412 ms), compatible with P3a, in response to a violation of local regularity. Violation of global regularity elicited an anterior negative response (228-586 ms), resembling the N400 component of ERPs. Importantly, the other indexes of auditory discrimination, such as MMN and P3b, were insignificant in ERPs to both regularity violations. The observed P3a and N400 components of ERPs may reflect prediction error signals in the implicit perception of sound patterns even if behavioral recognition was poor.

**Highlights:**

- Processing of hardly distinguishable harmonic tones differs from processing of complex patterns, which consist of these tones
- Implicit perception of local and global irregularities differs
- In passive listening, local irregularity elicits frontal positivity that is associated with an involuntary switch of attention
- In passive listening, global irregularity elicits late negativity that might reflect prediction error signals

## 1. Introduction

Predictive coding theory explains perception via a hierarchical model of prediction when the brain learns to predict sensory input by extracting regularities in the transition probabilities and detect unexpected deviants from these regularities (Friston, 2005). If the sensory input in the lower level does not match the prediction coming from higher levels, the prediction error adjusts the internal model, and further prediction is updated. As a result, the prediction model, with the most precise probability, determines our unconscious and conscious perception, which can be studied with event-related potentials (ERPs).

Several components of ERPs have been shown to reflect pre-attentive stimuli prediction. One of them is mismatch negativity (MMN), an index of auditory discrimination, proposed by R. Näätänen and his colleagues in 1978 (Näätänen et al., 1978). MMN is a component of the difference waveform of ERPs, which appears in response to rare (deviant) sounds comparing with ERPs to frequent (standard) sounds in odd-ball sequence, irrespective of participant’s attention to the stimuli (Auksztulewicz and Friston, 2015; Cacciaglia et al., 2019; Näätänen et al., 2001). MMN has been detected in the window 80-150 ms at frontocentral electrodes, and its amplitude directly correlated with the difference between stimuli (Näätänen et al., 2007). The increase of MMN amplitude was associated with more successful recognition of speech sounds (Näätänen & Alho, 1997) and learning (Winkler et al., 1999). MMN was diminished when subjects could not detect a difference between phonemes of a foreign language, but the MMN amplitude enhanced significantly with learning to recognize the foreign phonemes (Winkler et al., 1999). Later, these findings were consistently replicated in other studies (for review, Näätänen et al., 2007). Even though attention modulates MMN, it can be elicited without direct attention (Garrido et al., 2009), in light sleep stages (Atienza et al., 2002) and even in patients with disorders of consciousness (Morlet and Fischer, 2014).

In contrast to MMN, registered both in conscious and unconscious conditions, the P300 component of ERPs appears only when a participant is conscious and concentrates on the task (Sutton et al., 1967). Later, P300 was divided into two subcomponents: P3a and P3b. P3a was located at the frontocentral area in the time window from 150 to 250 ms, while P3b had temporal-parietal topography in the window from 300 to 600 ms. P3a is suggested to be a rebound of MMN reflecting the unconscious switch of attention to deviant stimuli (Ungan et al., 2019). While P3b is associated with conscious, direct attention as it was observed only in cases when subjects provided active responses supporting stimuli recognition (Polich, 2007). Therefore ERPs may reflect both unconscious and conscious levels of perception more precisely than only behavioral results (Atienza et al., 2002).

One experimental approach to study conscious versus unconscious perception was proposed by Bekinschtein et al. (2009) and named the local-global paradigm. The local-global paradigm helps to differentiate ERP indices of implicit (unconscious) recognition such as MMN and P3a, and explicit (conscious) recognition of stimuli such as P3b in clinical groups (Faugeras et al., 2012). Using MEG/EEG, it was found that local irregularity in the stimuli sequence elicited MMN and P3a even in non-communicating patients with disorders of consciousness (Bekinschtein et al., 2009; Faugeras et al., 2012; Shirazibeheshti et al., 2018) while global irregularity enhanced the P3b amplitude when a participant was conscious and actively engaged in the sound discrimination task (Bekinschtein et al., 2009; Chennu et al., 2013; Strauss et al., 2015).

Different components of ERPs appearing in response to either local or global irregularities can be explained from the combined point of view from the Predictive Coding model (Friston, 2005) and Global Neuronal Workspace Theory (Dehaene and Changeux, 2011). MMN and P300 have different brain sources that result in a distinct temporal span needed to infer the regularity. The local violation activates the primary auditory cortex (Bekinschtein et al., 2009). Using a mathematical model of Predictive Coding, Wacongne et al. (2012) revealed that MMN is an automatic response or low-level prediction error signal driven by transitional probabilities inferred from the close temporal neighborhood, as the MMN-like response to the local, short-term, irregularity was detected even in an unconscious state (Bekinschtein et al., 2009; Faugeras et al., 2011, 2012). P3a, as a rebound of MMN, had sources of generation in cingulate, frontal, and right parietal areas (Volpe et al., 2007). The more late P3b-like response to the global, long-term, irregularity was considered a high-level prediction error signal because it appeared only in case of direct attention (Bekinschtein et al., 2009; Strauss et al., 2015). According to the Global Neuronal Workspace Theory, P300 (P3b) is a conscious response driven by global stimulus properties inferred from a longer time range (Strauss et al., 2015). Moreover, the P3b response requires global network activation involving anterior cingulate, prefrontal, temporal, parietal, and occipital areas (Bekinschtein et al., 2009).

Importantly, unconscious (implicit) processing of stimuli from the environment underlies implicit learning, which is independent of directed attention and resources of declarative memory, unlike explicit learning (Yang and Li, 2012). New stimuli could appear ambiguous or hardly distinguishable; however, they still elicit the involuntary auditory response reflected in components of ERPs: N1-P2-N2 complex, MMN, and P3a (Nikjeh et al., 2009). These ERPs components are potential biomarkers of unconscious auditory perception reflecting mechanisms of implicit learning without a conscious understanding of the complex organization of stimuli patterns (Seger, 1994). If hardly distinguishable sounds are organized in sequences with long-time range irregularity temporal structure, possibly the discrimination of these sounds would transfer from implicit to explicit due to engagement higher levels of prediction (Anzulewicz et al., 2015; for review, Berlin, 2011 and Voss et al., 2012).

The current study investigated whether ERPs reflect an implicit perception of the sounds, which are difficult to differentiate for most healthy subjects. We hypothesized that sounds organized in patterns as in the passive local-global paradigm might facilitate transfer from the unconscious (implicit) level of stimulus recognition to the conscious level and simultaneously may elicit MMN and P3a (in local) and more pronounced P3a (in global regularity violation) compatible with active discrimination of deviant sounds in classical odd-ball paradigm. To test this hypothesis, we compared ERPs in response to hardly distinguishable non-verbal sounds in two conditions: active recognition of one deviant with the task to ignore the other deviant and standard sound, and passive listening of the same deviant and standard sounds presented in sequence with violation either local or global regularity.

## 2. Methods and materials

### 2.1. Participants and the study procedure

The study sample consisted of twenty healthy volunteers (mean age = 22.15 ± 2.4 years; 13 females) recruited for the project of memory reactivation during daytime nap (RFBR # 16-04-01403). Eligibility criteria to the participants: right-handed; aged 18-30 years; non-smokers; with no history of hearing impairments, brain injury, neurological diseases; non-ongoing users of medications. All participants were instructed to refrain from alcohol for 24 hours and from caffeine (i.e., tea and coffee) for 6 hours before the laboratory visit. The research protocols were compiled according to the Helsinki Declaration’s requirements, and the local ethical committee approved the study in the Institute of Higher Nervous Activity and Neurophysiology of the Russian Academy of Sciences (IHNA & NPh RAS). All participants were provided written informed consent and received a monetary reward (750 RUB).

We analyzed ERP and behavioral data obtained in the active three-stimulus odd-ball paradigm (OBP) and ERP data obtained in the local-global paradigm (LGP) during passive listening of sound sequences (Fig.1A).

**Fig1.**
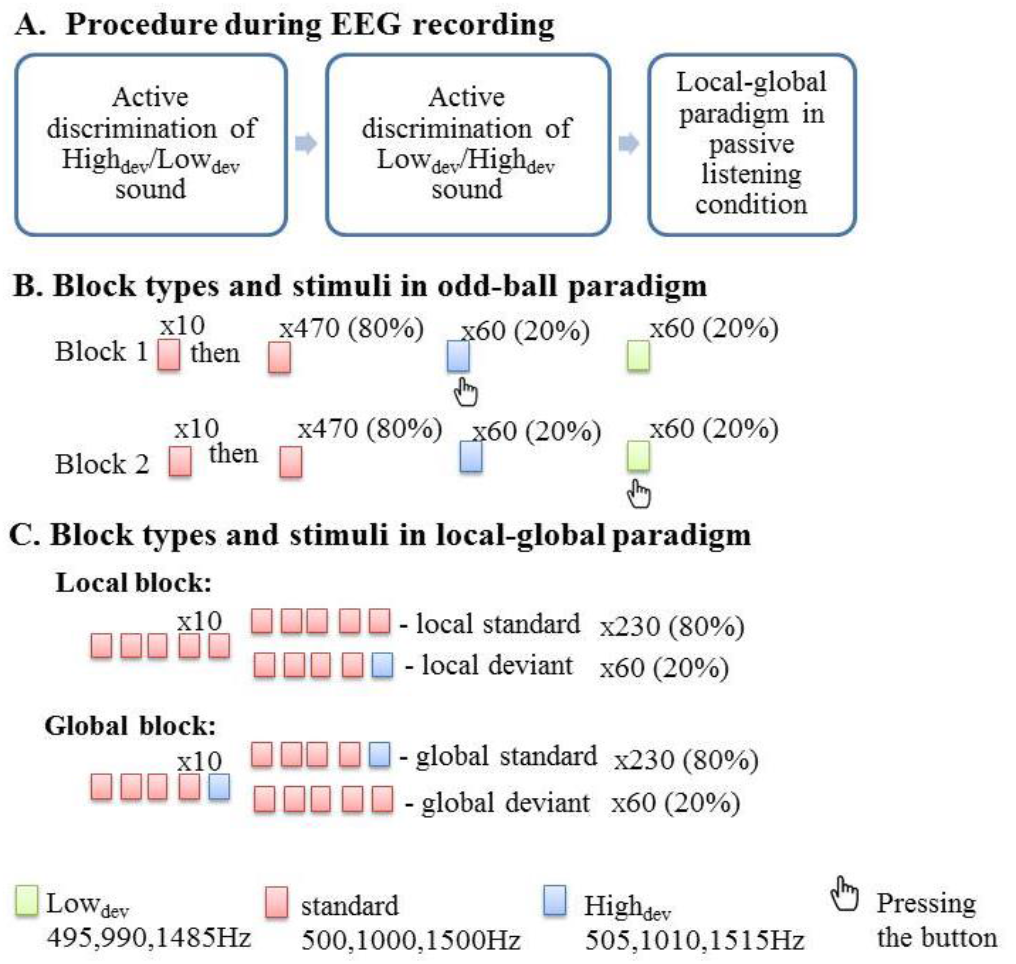
Experimental protocol. (A) EEG recordings were obtained during three sessions: during two active discrimination task and during perception of local and global blocks in passive listening condition. The goal was to evaluate whether the brain reacts to difficult to differentiate sounds organized in local and global irregularities in a passive state. (B) The first task was to see whether participants could differentiate the sounds. The participants were presented an active add-ball paradigm. During session 1 they pressed the button if they heard High_dev_, target deviant, they ignored Low_dev_, non-target deviant, and standard sound. During session 2 they pressed the button when they heard Low_dev_, target sound, and ignored High_dev_, non-target deviant, and standard sound. The trials were counterbalanced. (C) In a local block, frequent sequences with five identical sounds were termed “local standards”, and rare sequences in which the fifth sound was different were termed “local deviant”. Local mismatch effect is the difference between local deviant and local standard. In a global block, frequent sequences, in which the fifth sound was different, were termed “global standards”, and rare sequences with five identical sounds were “global deviants”. Global mismatch effect is the difference between global deviant and global standard. The participants were asked to think about nothing while local-global blocks were presented.

### 2.2. Auditory paradigms and stimuli

We used three types of sound stimuli with small differences in the frequency filling. In the active OBP, we presented two deviant stimuli and one standard stimulus (S) in the random order (Fig 1B). Standard was a harmonic tone composed of three sinusoidal partials of 500, 1000, and 1500 Hz. The deviant tones differed from standard tones in frequency as follows: (1) high deviant sound (High_dev_): 505, 1010, and 1515 Hz; (2) low deviant sound (Low_dev_): 495, 990, and 1485 Hz. In all three sounds, the intensity of the second and third partials was lower than the first partial by 3 and 6 dB. The duration of each sound was 75ms (including 5 ms rise and fall times), stimulus onset asynchrony (SOA) was 1000 ms with 50 ms jitter. The stimuli were presented via headphones at 60 dB above the individual subject’s hearing threshold with equal phase and intensity at both ears. Standards appeared with 80% frequency, and High_dev_ and Low_dev_ were presented with an equal frequency of 10%. Altogether 480 standard and 120 deviant sounds (60 - High_dev_, 60 - Low_dev_) were presented in 10 min 75 s block, where the first ten trials always were standards. In one block of OBP, we asked participants to press a mouse button when they heard High_dev_ – target deviant – and ignore other sounds (Low_dev_ – non-target deviant – and Standards). In another block, we asked to respond to Low_dev_ and ignore other sounds (High_dev_ and Standards). The order of blocks was assigned randomly between subjects.

In the passive listening condition, we used LGP adapted from the study of Bekinschtein et al. (2009) (Fig.1C). The paradigm consisted of two blocks with a violation of local and global regularities. The first four stimuli in the trial were identical, while the fifth stimulus was either the same or different. Local irregularity block contained 80% local standard (LS) SSSSS trials and 20% local deviant (LD) SSSSD trials; both appeared in the random order. The global irregularity block contained 80% global standard (GS) SSSSD trials and 20% global deviant (GD) SSSSS trials. Standard was a harmonic tone composed of three sinusoidal partials of 500, 1000, and 1500 Hz. In 14 subjects (Group 1), Deviant was High_dev_, 505, 1010, and 1515 Hz, and in 6 people (Group 2) - Low_dev_, 495, 990, and 1485 Hz. The duration of the sounds was 50 ms (including 5 ms rise and fall times). Each sound in the trial sequence was separated by 100 ms interstimulus interval (ISI), while SOA between trials was 1400 ± 50 ms. The stimuli were presented via headphones 40 dB during LGP above the individual subject’s hearing threshold with equal phase and intensity at both ears. The presentation of GLP sounds consisted of four blocks (2 with local and 2 blocks with global irregularity). 125 standard and 30 deviant trials were presented in 4 min 13 s block; the first ten patterns were standard one.

Presentation of stimuli sequences, synchronization with EEG recording and registration of subjects’ responses was done by E-Prime 1.2 («Psychology Software Tools,» Pittsburgh, PA, US).

### 2.3. Data acquisition

Continuous EEG data were acquired by a digital 19-channel EEG amplifier Encephalan-EEGR-19/26 (Medicom, Taganrog, Russia), where 19 AgCl electrodes (Fp1, Fp2, F7, F3, Fz, F4, F8, T3, C3, Cz, C4, T4, T5, P3, Pz, P4, T6, O1, O2) were placed according to an extended international 10–20 system. Additional 4 electrodes were used for acquisition of electrooculograms (EOG). For recording vertical EOG, electrodes were placed 1 cm above and below the left eye. For recording horizontal EOG, electrodes were placed 1 cm lateral from the outer canthi of both eyes. Left and right mastoid electrodes served as reference channels for the monopolar design of EEG recording. Impedance was kept below 10 kΩ. A band-pass filter was set from 0.05 to 70 Hz, and the sampling rate was 250 Hz.

### 2.4. EEG data preprocessing

We analyzed data in BrainVision Analyzer 2.0 (Brain Products GmbH, Gilching, Germany). First, we applied a high pass filter of 0.5 Hz. To remove eye-movement and blink-related EEG artifacts, we conducted Independent Component Analysis (ICA). Then we applied low-pass filter 30 Hz and a notch filter at 50 Hz. After that, we visually inspected and manually cleared continuous EEG from periods with muscle and motion artifacts. In OBP, the epochs were segmented from 200 ms before to 600 ms after the onset of the sound. In LGP, epochs were segmented from 800 ms before to 600 ms after the onset of the fifth sound in the sound trial. We used baseline correction −200 to 0 ms window before the presentation of the fifth tone.

Additionally, we performed an automatic artifact rejection of EEG epochs. We defined the following criteria for artifact rejection: a voltage step of more than 100 μV/ms a voltage difference of less than – 100 μV or more than 100 μV and a maximum voltage difference of more than 100 μV within 100 ms intervals. The artifact-free epochs were averaged separately in response to standards, High_dev_, and Low_dev_ in OBP, and in response to the fifth sound in LD, LS, GD and GS trials in LGP. In both OBP and LGP, the first 10 standard sounds were not included in the averaging.

### 2.5. Statistical analyses of behavioral and ERP data

To examine behavioral task performance in the sound discrimination task, we considered reaction times (RTs), hit rates (hits), d-prime sensitivity index (d’), and decision criterion (C) (Macmillan and Creelman, 1991). These variables were subjected to the Mann–Whitney U test in order to compare performance between groups. The latencies of ERPs peaks at different electrodes were subjected to the Wilcoxon signed-rank test. Spearman’s rank correlation tested the association between RTs and ERPs latency. Statistical comparison of ERPs was performed by cluster permutation tests corrected for multiple comparisons over time (Fieldtrip, Monte Carlo method, 500 permutations, the alpha level was 0.05) in the window between 0 and 600 ms after the stimulus onset in OBP, and after the fifth stimulus onset in the trial for LGP.

## 3. Results

### 3.1. Behavioral and ERP data in the active three-stimulus odd-ball paradigm

Group 1 (N=14) and Group 2 (N=6) differed in d’ (p=0.063, Z=-1.856 for hits; p=0.058, Z=1.898 for false alarms, p=0.023, Z=-2.269 for d’; and p=0.65, Z=0.454 for C). This difference may be explained by a small number of subjects in Group 2; thus, we combined both groups’ results. In general, the accuracy in discrimination of sounds was low (Average hits = 0.40 ± 0.21, false alarms = 0.11 ± 0.13, d’ = 0.85 ± 0.7, and C = 1.35±0.61). The significant deviation of the d’ from 0 indicates participants’ ability to reliably discriminate deviants from noise at the group level. However, d’ index lower than 2 and the substantial standard deviation (Fig.2) in the group indicate difficulties in making an unambiguous decision in discrimination of sounds (Macmillan and Creelman, 1991). This assumption is also supported by numerous incorrect choices of standard sounds instead of target deviants (66 %).

**Fig.2.**
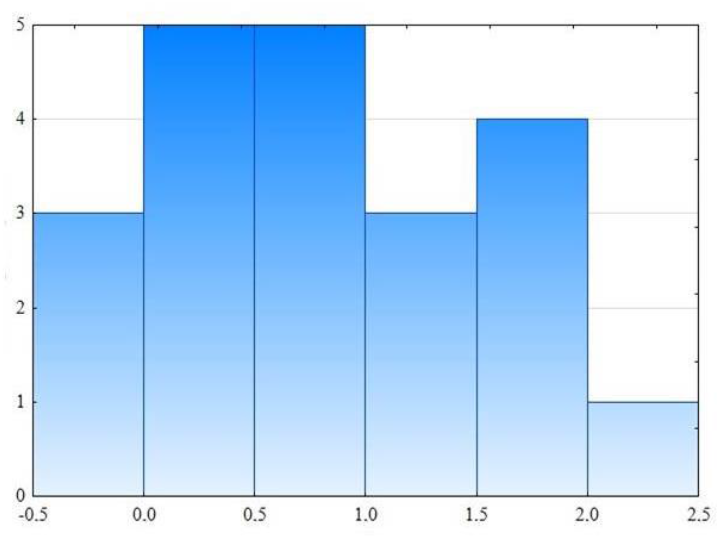
The histogram on d’ as the accuracy in discrimination of sounds (N=20). X-axis – d’; Y-axis – number of participants.

To check whether there is a difference between subjects who have differentiated sounds better and those who have not, using cluster-level statistics, we compared ERPs to targeted deviant sounds (required active discrimination) and ERPs to standards of two equal groups (N=10 in both groups) divided according to the median of d’ results (median = 0.86, p = 0.005). There was no difference in ERPs between groups.

Uniting these two groups, we found differences between ERPs to deviant (when this deviant was either target or non-target one) and ERPs to standard. Non-target deviant elicited frontal negativity (100-132 ms, p = 0.03) (Fig.3A). The targeted deviant elicited a more prominent posterior negativity peaked around 500 ms (456-596 ms, p = 0.008) (Fig.3B). The deviant differed from standard only in amplitude. There was no difference in latency between standard and deviant sounds at Cz and Pz electrodes was (p = 0.756, Z = 0.31 and p = 0.074, Z = 1.784, respectively). To check whether this posterior negativity reflects a preparation to motor response, we correlated the latency of this negative peak for both deviant and standard sounds in 456-596 ms window with RTs (541 ± 156 ms and 517 ± 131 ms average RTs of deviant and standard, respectively). No correlation was found between them. The only significant correlation was between RTs of deviant and RTs of standard: the longer it took time to process standard, the longer deviant latency was (r=0.647).

**Fig3.**
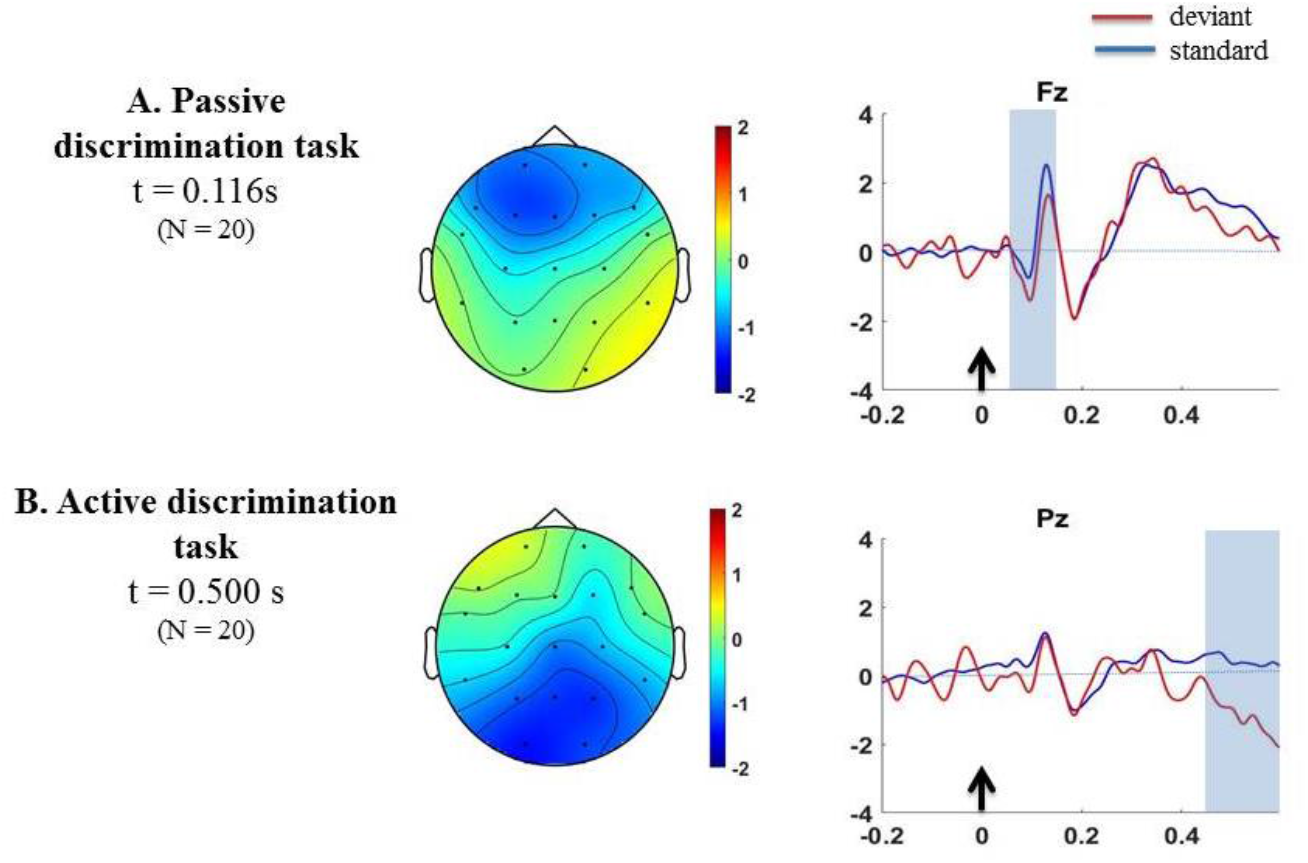
Event-related responses in odd-ball paradigm in a passive listening condition. MMR in EEG. (A) Topography of the mismatch response (non-target deviants minus non-target standards) is shown at t = 114 ms after the fifth sound presentation. Cluster based significant effect (p<0.05) in 100–to–128-ms time window is represented by blue shaded region. (B) The topography of the mismatch response (target deviants minus non-target standards) is shown at t = 500 ms after the fifth sound presentation. Cluster based significant effect (p<0.05) in 456–to–598-ms time window is represented by blue shaded region. The black arrow is a sound either standard or deviant. X-axis – time in seconds; Y-axis – amplitude in microvolts.

### 3.2. ERPs to local and global irregularities in passive listening condition

ERPs to violations of short-term regularities were analyzed by comparing responses to LD and LS as contrast termed the local effect. A significant local effect was found using cluster-level statistics at around 350 ms, reflecting in the broad anterior positivity, comparable to P3a wave (284-412 ms, p = 0.002), in response to LD (Fig.4A).

**Fig4.**
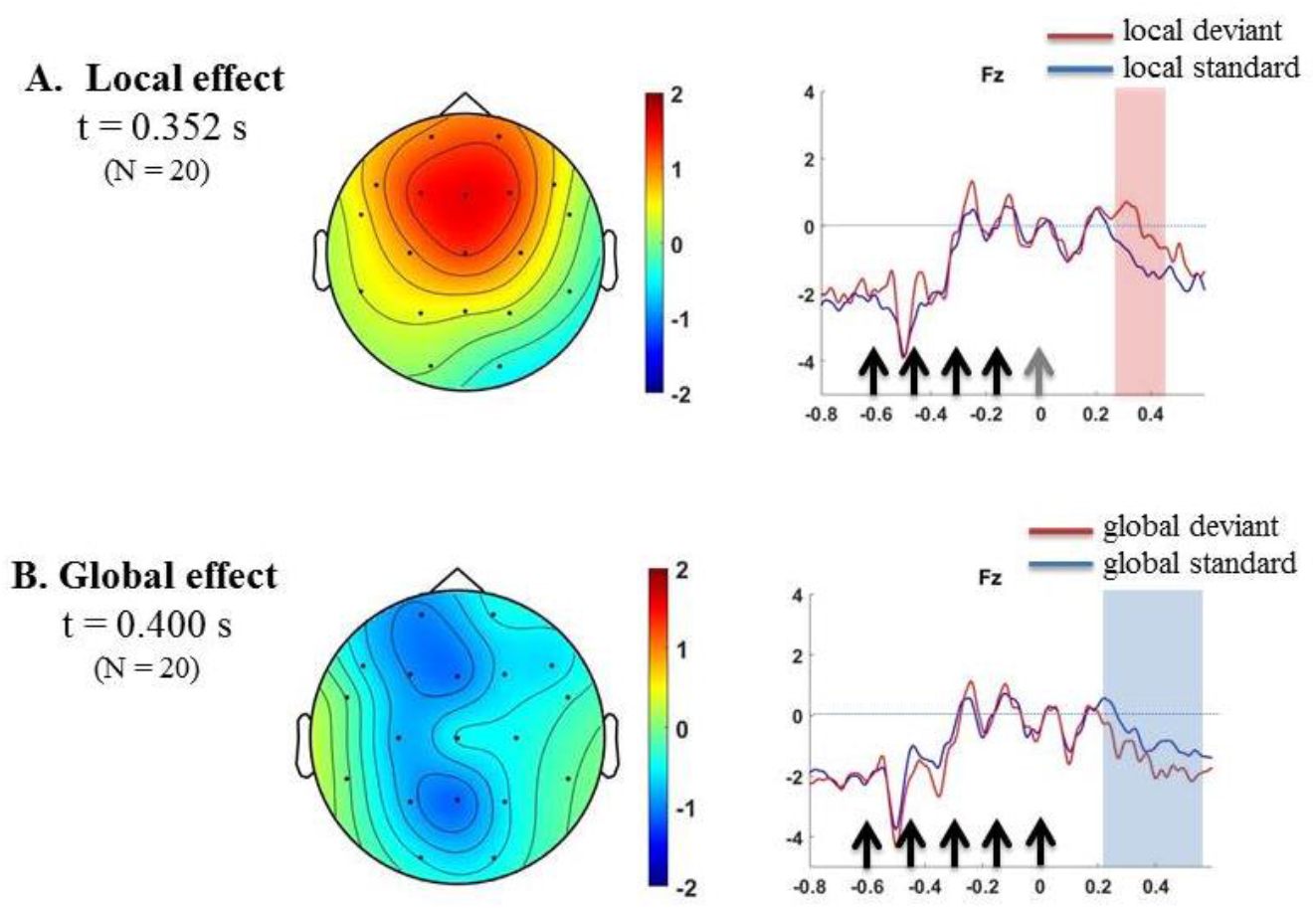
Event-related responses to local and global irregularities in passive listening condition. (A) Local mismatch response in EEG. The topography of the local effect (local deviants – local standards) is shown at t = 350 ms after the fifth sound. Cluster based significant effects (p<0.05) in 284 – to – 412 ms time window are represented by pink shaded region. Black arrows are standard sounds, a grey arrow is a deviant sound. (B) Global MMR in EEG. The topography of the global effect (global deviants – global standards) is shown at t = 400 ms after the fifth sound. Cluster based significant effects (p<0.05) in 228-to-586 ms time window are represented by blue shaded region. Black arrows are standard sounds. X-axis – time in seconds; Y-axis – amplitude in microvolts.

Due to the global effect, ERPs to violations of long-term regularities were analyzed by comparing responses to GD and GS. A significant global effect was observed at around 400 ms, represented by more prominent anterior negativity in response to GD, comparable to the N400 wave (228-586 ms, p = 0.04) (Fig.4B).

## 4. Discussion

The current study investigated whether ERPs reflect an implicit perception of the sounds, which are difficult to differentiate for adult subjects without hearing deficit. We hypothesized that even in passive listening, sounds organized in patterns as in the local-global paradigm would elicit ERPs, such as MMN and P3a for local irregularity, and more pronounced P3a for global irregularity.

According to signal detection theory, participants could differentiate stimuli; however, the averaged recognition reflected in d’ sensitivity was below 1, supporting the difficulty of correct choice for participants (Macmillan and Creelman, 1991). Probably due to this difficulty of the explicit choice of “correct” deviant, we have registered neither significant MMN nor the cognitive component P300 in response to the targeted (correct) deviant in the active discrimination task. We only observed a significant increase in the negative amplitude of ERPs in response to non-target (incorrect) deviant compared with ERPs to standard in the time window 100-132 ms, which could be associated with early MMN. Our data are consistent with previous results showed that the amplitude of MMN directly correlates with behavioral performance (Näätänen and Alho, 1997; Ukraintseva et al., 2018) and can be modified by the top-down influence of selective attention depending on task goals (Oades and Dittmann-Balcar, 1995; Sussman et al., 2002). Instead of MMN and P300, we detected the late negative posterior-occipital potential around 500 mas after stimulus onset in response to targeted deviants. Based on its latency and localization, this negativity resembles a readiness potential to the presented stimuli (Brunia et al., 2011). However, in our study, the rate of false alarms (a reaction to wrong non-targeted deviant, or choosing standard sound) was also higher (66 %), resulting in a considerably low d’ index of sound detection (0.85). In this case, we should observe almost equal readiness potentials to correct, incorrect deviants, and standards. An appearance of readiness potential only in response to correct-to-choose deviant suggests that this late negativity is more likely associated with the implicit perception of stimuli than with the readiness to perform a motor act after sound discrimination. Additionally, a correlation analysis did not show a significant relationship between reaction times and latencies of this negative component.

We did not observe significant MMN and P300 in the active discrimination task, supporting the behavioral data on ambiguous sound recognition. Further, we hypothesized that sounds organized in patterns as in the passive local-global paradigm might induce a more pronounced effect on the implicit sound recognition, and correspondingly on the ERP indexes of pre-attentive auditory discrimination. To check this, we separately investigated ERPs in response to sound sequences with local and global irregularities. We did not detect a statistically significant MMN on the local effect but observed frontocentral positivity comparable to the P3a component according to its latency and scalp localization. The P3a is a subcomponent of P300 associated with unconscious exogenous attention (Polich, 2007). This component was elicited in conscious, healthy individuals, and patients with disorders of consciousness (Faugeras et al., 2012). In our study, the P3a component elicited in response to the violation of local regularity in the sequence of hardly distinguished sounds, which were also presented in the preceding active discrimination task. Thus, the observed P3a may reflect the unconscious detection of a deviant stimulus that appeared as local irregularity in the sound sequence.

As we expected, we did not detect the late positive parietal-occipital subcomponent P3b reflecting the global effect in the passive listening condition. The absence of P3b in our study is consistent with previous results showing that P3b requires direct attention (Bekinschtein et al., 2009; Faugeras et al., 2012; Strauss et al., 2015). P3b is elicited only when the stimulus is consciously recognized using a strategic process necessary to detect and count different patterns (Bekinschtein et al., 2009). Notably, that in response to the global irregularity, we observed significant anterior negativity at latencies of the N400 component of ERPs. The language studies frequently reported an association of N400 with pre-attentive semantic processing (Erlbeck et al., 2014; Kiefer, 2002; Schneider and Shiffrin, 1977). The semantic N400 was usually more prominent at the central and parietal electrodes (Rohaut and Naccache, 2017). Predictive coding theory explains semantic N400, pre-attentive language comprehension, in terms of probabilistic information and categorical template matching (Bornkessel-Schlesewsky and Schlesewsky, 2019; Grisoni et al., 2017; Vespignani et al., 2010). Our findings instead reflect the appearance of non-semantic N400, having a frontocentral topography and recorded during the detection of non-verbal “semantic” patterns (Cummings et al., 2006). M. Kutas and K.D. Federmeier (2011) also described a frontal N400 (FN400) registered as a response to meaningful non-speech stimuli in a recognition task. A similar N400 component was reported in implicit learning (Rabovsky et al., 2018), which was the opposite of familiarity in the awareness of memory retrieval and explicit learning (Voss and Paller, 2009). Also, the observed late negative component of ERPs in response to global irregularity can also be associated with late discriminative negativity (LDN) (Čeponienė et al., 1998; Cheour et al., 2001) characterized by anterior topography and late window 300-500 ms in children studies (Strotseva-Feinschmidt et al., 2015). Even though the function of LDN is still unclear, it might be an index of the automatic but higher-order discrimination process of complex auditory stimuli (Horváth et al., 2009), which based on the previous finding on N400 or late negativity, we may only assume that the observed late negativity in both OBP with active discrimination task and global irregularity in passive listening of sounds in LGP reflects the second hierarchic level of implicit error prediction according to the predictive coding theory (Lecaignard et al., 2015). However, further research is needed to explore the neural mechanisms of implicit processing of complex auditory sequences.

## 5. Conclusions

Our study shows that active detection of hardly distinguishable sounds in an odd-ball paradigm elicited late posterior negativity (around 500 ms). When these sounds were organized in patterns with local or global violations, local irregularity elicited P3a, possibly associated with an involuntary switch of attention toward the deviant pattern. In contrast, global irregularity elicited late negativity (around 400 ms), which might reflect prediction error signals in the implicit perception of sound patterns even if behavioral recognition was poor.

## Data availability statement

The datasets generated for this study are available on request to the corresponding author.

## Author contribution

KL, YuU conducted the experimental tasks and analyzed data. KL and OM wrote the manuscript.

## Acknowledgments

We gratefully acknowledge the help of the volunteers who participated in this study. We thank our colleagues, especially Olga Kashevarova and Beatriz Bermúdez-Margaretto for technical support, Alexandra Polishchuk, Mikhail Shilov, and Kaushik Sake for help in conducting the experiments as well as Yury Shtyrov and Andriy Myachykov for insightful comments on an earlier draft of the manuscript.

